# Necessary Role of Acute Ceramide Formation in The Human Microvascular Endothelium During Health and Disease

**DOI:** 10.1101/2023.06.02.543341

**Authors:** Gopika SenthilKumar, Boran Katunaric, Zachary Zirgibel, Brian Lindemer, Maria J. Jaramillo-Torres, Henry Bordas-Murphy, Mary E. Schulz, Paul J. Pearson, Julie K. Freed

## Abstract

**Background:** Elevated plasma ceramides independently predict adverse cardiac events and we have previously shown that exposure to exogenous ceramide induces microvascular endothelial dysfunction in arterioles from otherwise healthy adults (0-1 risk factors for heart disease). However, evidence also suggests that activation of the shear-sensitive, ceramide forming enzyme neutral sphingomyelinase (NSmase) enhances vasoprotective nitric oxide (NO) production. Here we explore a novel hypothesis that acute ceramide formation through NSmase is necessary for maintaining NO signaling within the human microvascular endothelium. We further define the mechanism through which ceramide exerts beneficial effects and discern key mechanistic differences between arterioles from otherwise healthy adults and patients with coronary artery disease (CAD).

**Methods:** Human arterioles were dissected from otherwise discarded surgical adipose tissue (n=123), and vascular reactivity to flow and C2-ceramide was assessed. Shear-induced NO production was measured in arterioles using fluorescence microscopy. Hydrogen peroxide (H_2_O_2_) fluorescence was assessed in isolated human umbilical vein endothelial cells.

**Results:** Inhibition of NSmase in arterioles from otherwise healthy adults induced a switch from NO to H_2_O_2_-mediated flow-induced dilation within 30 minutes. In endothelial cells, NSmase inhibition acutely increased H_2_O_2_ production. Endothelial dysfunction in both models was prevented by treatment with C2-ceramide, S1P, and an agonist of S1P-receptor 1 (S1PR1), while the inhibition of S1P/S1PR1 signaling axis induced endothelial dysfunction. Ceramide increased NO production in arterioles from healthy adults, an effect that was diminished with inhibition of S1P/S1PR1/S1PR3 signaling. In arterioles from patients with CAD, inhibition of NSmase impaired dilation to flow. This effect was not restored with exogenous S1P. Although, inhibition of S1P/S1PR3 signaling impaired normal dilation to flow. Acute ceramide administration to arterioles from patients with CAD also promoted H_2_O_2_ as opposed to NO production, an effect dependent on S1PR3 signaling.

**Conclusion:** These data suggest that despite key differences in downstream signaling between health and disease, acute NSmase-mediated ceramide formation and its subsequent conversion to S1P is necessary for proper functioning of the human microvascular endothelium. As such, therapeutic strategies that aim to significantly lower ceramide formation may prove detrimental to the microvasculature.

## NOVELTY AND SIGNIFICANCE

### What is known?

- Ceramides are elevated in patients with endothelial dysfunction, independently predict major adverse cardiac events, and promote microvascular endothelial dysfunction in arterioles isolated from otherwise healthy adults.
- An important source of intracellular ceramide formation is via neutral sphingomyelinase (NSmase) enzyme, inhibition of which reduces plaque progression and vascular dysfunction in preclinical models.

### What New Information Does This Article Contribute?

- Acute neutral-sphingomyelinase mediated ceramide formation is necessary for maintaining nitric oxide (NO) mediated flow-induced dilation of arterioles from otherwise healthy adults.
- Ceramide exerts beneficial effects on the endothelium through its conversion to sphingosine-1-phosphate (S1P) and activation of S1P-receptor 1 and 3.
- Ceramide formation is necessary for dilation to flow in arterioles from patients with CAD. However, ceramide primarily signals through S1PR3 in these arterioles, resulting in generation of hydrogen peroxide as opposed to NO.

Cardiovascular disease (CVD) remains a leading contributor to morbidity and mortality worldwide. Emerging evidence highlights microvascular function as a key, early predictor of CVD outcomes. However, there are currently no targeted strategies for improving microvascular function. We and others have shown that ceramide is detrimental to the microcirculation, and as such therapeutics that inhibit ceramides formation are emerging as an attractive strategy to mitigate CVD risk. However, this study using *human* microvessels, highlights that ceramide formation is necessary for proper microvascular endothelial functioning during both health and disease. As such, therapeutics that aim to significantly lower ceramide levels, especially through inhibition of neutral sphingomyelinase enzyme, may actually prove detrimental to the vasculature. Furthermore, we mechanistically show that ceramide signals through S1P in both health and disease, although ceramide produces vasoprotective NO during health and pro-inflammatory hydrogen peroxide in arterioles from patients with CAD. This work not only highlights key differences in sphingolipid signaling between health and disease, but also suggests that further understanding of downstream S1P signaling may provide novel avenues for therapeutic targeting. Additionally, the mechanisms described in this work highlight key differences in sphingolipid signaling between humans and previously published preclinical studies, which have important implications for clinical translation.

## INTRODUCTION

The primary role of the microvasculature is to match organ perfusion with metabolism through finely tuned adjustments in vascular tone. To make such adjustments, an arteriole must sense an increase in blood flow (demand) and respond by increasing its diameter (flow-induced dilation; FID). This important physiological response relies on a functioning endothelium to generate compounds capable of causing relaxation of the underlying vascular smooth muscle. In microvessels from healthy patients, FID is mediated by endothelial generation of the anti-inflammatory, anti-thrombotic compound nitric oxide (NO)^1^. Endothelial dysfunction, or the inability of the endothelium to release NO in response to shear, is observed in both conduit arteries and microvessels from patients diagnosed with or possessing multiple risk factors for coronary artery disease (CAD)^1–3^. While conduit arteries experience a loss of dilator capacity in response to flow^4^, resistance arterioles can compensate for the loss of NO by releasing hydrogen peroxide (H_2_O_2_)^5^ to elicit dilation. While this transition in mechanism allows for dilation during disease, H_2_O_2_ release promotes a pro-inflammatory, pro-thrombotic, and pro-atherosclerotic perivascular environment^6, 7^.

We have previously shown that chronic exposure to ceramide, a sphingolipid elevated in individuals with endothelial dysfunction^8^ and increases risk of major adverse cardiac events (MACE) in otherwise healthy individuals^9^, promotes microvascular endothelial dysfunction in arterioles isolated from nonCAD patients (having 1 or less risk factors for CAD)^10^. Suppressing ceramide formation or increasing ceramide metabolism restores NO-mediated FID in arterioles from patients with CAD^10–12^. These data are in agreement with preclinical studies highlighting the detrimental effects of ceramide on the vasculature^13–15^ and therapeutics that aim to decrease ceramides are emerging as an attractive strategy to mitigate future cardiac disease risk^12, 16, 17^.

An important source of intracellular ceramide is via metabolism of plasma membrane sphingomyelin by neutral sphingomyelinase (NSmase), an enzyme rapidly activated by shear stress^18^ to increase intracellular ceramide levels (in <1min). Inhibition of NSmase reduces atherosclerotic plaque progression^13^ and prevents western-diet induced vascular dysfunction^14^ in preclinical models. However, ceramide has also been identified as a key signaling lipid within the endothelium and NSmase activity has been linked to activation of downstream targets that favor NO production, such as endothelial nitric oxide synthase (eNOS)^19, 20^. In addition, metabolites of ceramide, such as Sphingosine-1-phosphate (S1P), have also been shown to stimulate production of NO through the activation of S1P receptor 1 (S1PR1)^21^.

Here, we explore a unifying theory to explain the seemingly paradoxical observation that ceramide is capable of having both beneficial and pathogenic roles in microvascular function. We hypothesized that ceramide formation through NSmase is necessary for maintaining a healthy, NO-producing microvascular endothelium. We further postulate that this source of ceramide metabolizes to S1P which subsequently activates S1PR1 to increase NO levels. Using resistance arterioles isolated from patients, we also were able to delineate differences in ceramide/S1P signaling in those diagnosed with CAD versus those with only 0-1 risk factors for heart disease. To our knowledge, this is the first study that addresses a critical knowledge gap in our understanding of how ceramide, a lipid that if accumulated intracellularly and within plasma increases one’s cardiovascular risk, is also a necessary mechanistic component of NO signaling within the human microvasculature.

## METHODS

### Human Tissue Collection

Fresh surgical, otherwise discarded adipose tissue (visceral, peritoneal, or subcutaneous) were acquired and placed in ice-cold HEPES buffer. When possible, patients scheduled to undergo elective procedures that commonly result in discarded adipose tissue were consented prior to surgery. Otherwise, discarded tissue was collected through the Medical College of Wisconsin tissue bank along with de-identified information on sex, age, race, CAD risk factors and related medications. Patients’ de-identified information was collected using the Medical College of Wisconsin REDCap database. Patients that had 0-1 risk factor(s) for coronary artery disease (hyperlipidemia, congestive heart failure, diabetes mellitus type 1 or 2, hypertension, active smoker) were classified as ‘healthy’ (nonCAD), and patients with a clinical diagnosis of CAD based on percutaneous coronary intervention were classified as CAD. Tissue from patients with a positive COVID-19 diagnosis within the past 6-months were excluded. All protocols were approved by the Institutional Review Board at the Medical College of Wisconsin.

### Human Microvascular Function Studies

Resistance vessels (mean±SD; 109.03±39.84μm) were isolated from adipose tissue, cannulated on glass micropipettes of matched impedance within an organ chamber, and maintained in pH and temperature (37°C) controlled KREBs buffer. The chamber was aerated (74% N2/21% O2/5% CO2 mixture) and micropipettes were connected to fluid reservoirs containing KREBs buffer to allow for pressurization. Vessels were equilibrated for 30 min at 30mmHg, followed by 60mmHg for 30 min and exposed to various treatments (Supplemental Table 2). Arterioles were pre-constricted to 30-70% of their passive intraluminal diameters with endothelin-1 (ET-1; 0.2-2nM). Vessels that maintained tone (<5% change in vessel diameter for 10 min) were subject to further experimentation. To induce flow, the reservoirs were moved in equal and opposite direction, and changes in internal luminal diameter following 5-min exposure to graded changes in pressure gradient (5–100 cmH2O; index of flow) was measured via videomicroscopy. This protocol was originally described by Kuo et al^22^., and has been extensively used in our laboratory^11, 23, 24^. Following the flow experiment, vessels were treated with 100µM papaverine (endothelial-independent dilator) to determine the vessel’s maximal diameter and assess smooth muscle function. Vessels that dilated less than 70% of their maximal passive diameters following papaverine treatment were excluded. Percent change in intraluminal diameter at each pressure gradient was calculated with 0% representing the endothelin-1 pre-constricted diameter and 100% representing the vessel’s maximal diameter. No more than 3 experiments were conducted per vessel.

### Agonist-Induced Dilation of Human Microvessels

Human arterioles were isolated, cannulated, and pressurized as previously described, and exposed to increasing doses (10^-9^ to 10^-5^ M) of C2-ceramide, dihydroceramide, or vehicle control (dimethyl sulfoxide vol/vol) in 1-min intervals. Changes in internal arteriolar diameter were recorded via videomicroscopy. Vessels were then treated for 30 minutes at 60mmHg with various inhibitors (Supplemental Table 2) prior to being exposed to increasing doses of ceramide (10^-9^ to 10^-5^). Vessels were then treated with 100µM papaverine, and vessels that dilated less than 70% of their maximal passive diameters following papaverine administration were excluded. Dilation with increasing ceramide doses was calculated as percent of maximal dilation.

### Measurement of Nitric Oxide Formation

Isolated arterioles from nonCAD patients were cannulated and intraluminally infused with the Enzo Life Sciences Nitric Oxide probe (Enzo Life Sciences, Enzo Biochem, Farmingdale, New York; 1:400 in HEPES buffer). Vessels were exposed to the probe for 1 hr at 30mmHg pressure, followed by 30 min at 60mmHg during which treatments were added. Images were acquired at 10x using the Olympus IX73 microscope (650/670nm excitation/emission). Following the acquisition of a baseline fluorescence image, vessels were exposed to 100cmH2O pressure gradient (maximal flow) and images were acquired 1 min and 5 min following flow exposure.

Vessels were washed 3 times with HEPES buffer and exposed to a second treatment group with 30 min incubation. Probe specificity for NO was confirmed using the nitric oxide synthase inhibitor L-NAME (100µM). No more than 3 experiments were conducted per vessel. Image J software was used to outline the vessel and total corrected fluorescence [integrated density – (vessel area*mean background intensity)] was measured per image^25^. Percent change in total corrected fluorescence from baseline is reported.

### Cell Culture

Pooled human umbilical vein endothelial cells (HUVECs; ATCC Manassas, VA, USA) were cultured in 5% fetal bovine serum containing endothelial cell growth medium (EGM-2; Lonza, Basel, Switzerland). Media was supplemented with heparin, hydrocortisone, ascorbic acid, and growth factors (EGM-2 SingleQuot; Lonza, Basel, Switzerland). Cells were used at passage 2-5 and plated onto 8 well chamber slides (Ibidi, Gräfelfing, Germany) at 70-80% confluency.

### Measurement of H_2_O_2_ in Cultured Endothelial Cells

Bright field images were first obtained at 20x to select imaging fields across all wells (3-4 fields per well). The peroxy yellow 1 (PY1) probe (1:400 in live cell imaging solution) with different treatments was then added to each well to measure H_2_O_2_ production over time. A baseline fluorescence image was acquired, and then images were acquired at 60-, 90-, and 120-minutes using a Keyence fluorescence microscope (1/30 second exposure; 488/510nm excitation/emission). Images were analyzed by a technician blinded to treatment. Mean fluorescence intensity was calculated for all images using Image J. Percent change in intensity from baseline per image was calculated and averaged per well. Signal specificity for H_2_O_2_ was confirmed with the addition of pegylated-catalase (catalase, 500 U/mL).

### Statistical Analysis

Patient characteristics between health and disease was compared using a two-tailed t-test for continuous variables and chi-square analysis for categorical variables. To assess whether overall graded response to flow varied between untreated and treated vessels, a non-linear regression was performed with a 4-parameter fit. The minimum was constrained to 0 and the maximum was constrained to 100, and the EC50 was constrained to be greater than 0. Best fit lines are represented as dotted lines on graphs, and 95% confidence interval of EC50s are reported in results. To assess the mediator (NO vs H_2_O_2_) of dilation, we evaluated whether overall dilation to flow/ceramide doses was impaired in presence of treatment+inhibitor (L-NAME or catalase) using a two-way repeated-measures ANOVA with pressure gradients/ceramide doses and treatments as parameters. For comparing changes in NO and H_2_O_2_ fluorescence intensity over time, a two-way repeated measures ANOVA was used with treatment and time as parameters.

When multiple comparisons were made, Holm-Sidaak multiple comparisons test was performed. The results based on ANOVA analysis are reported as difference between mean (DBMs) %±SE of treatment groups, with p-values for overall treatment vs treatment+inhibitor effects (treatment effects), as a function of time/dose (interaction effect), or at a particular time/dose (multiple comparisons test). Significance was defined as p<0.05. All analyses were performed using GraphPad Prism, v.9.1.2.

## RESULTS

Samples from 123 patients (91 healthy, 32 CAD) were included. Average age of healthy patients was 44.2±12.3 years and CAD patients was 65.3±9.7 years (p<0.001). 91% (83/91) of healthy patients and 38% (12/32) of CAD patients were female (p<0.001). BMI and race/ethnicity did not vary between groups. Other demographic information is detailed in Supplemental Table 1.

### Inhibition of neutral sphingomyelinase in nonCAD arterioles impairs NO-mediated FID

To evaluate whether inhibition of NSmase impairs NO-mediated FID of nonCAD human arterioles, microvessels were treated with GW4869 (4μM, 30min) and exposed to increasing flow rates. Compared to control arterioles, inhibition of NSmase did not affect overall maximal dilation to flow however triggered an amplified dilatory effect at lower flow rates (Figure 1A; 95% CI EC50 of best fit curves for control 15.74-37.34, n=9 vs GW4869, n=5: 2.73-8.76, p<0.001; Figure 1A). Treatment with vehicle alone did not impair FID compared to control (Supplemental Figure 1A). FID was maintained in the presence of the NOS-inhibitor L-NAME during NSmase inhibition (DBM GW4869, n=5 vs GW4869+L-NAME, n=4; -19.22±12.38; p=0.165, Figure 1B). The presence of L-NAME also eliminated the heightened dilatory effect to flow at lower flower rates that was observed with GW4869 alone (Figure 1C; 95% CI EC50 of best fit curves for control, n=9 and GW4869+L-NAME, n=4: 16.04-31.19, p=0.649). NO-mediated dilation was preserved in arterioles treated with vehicle control as FID was diminished in the presence of L-NAME (Supplemental Figure 1B).

**Figure 1.**
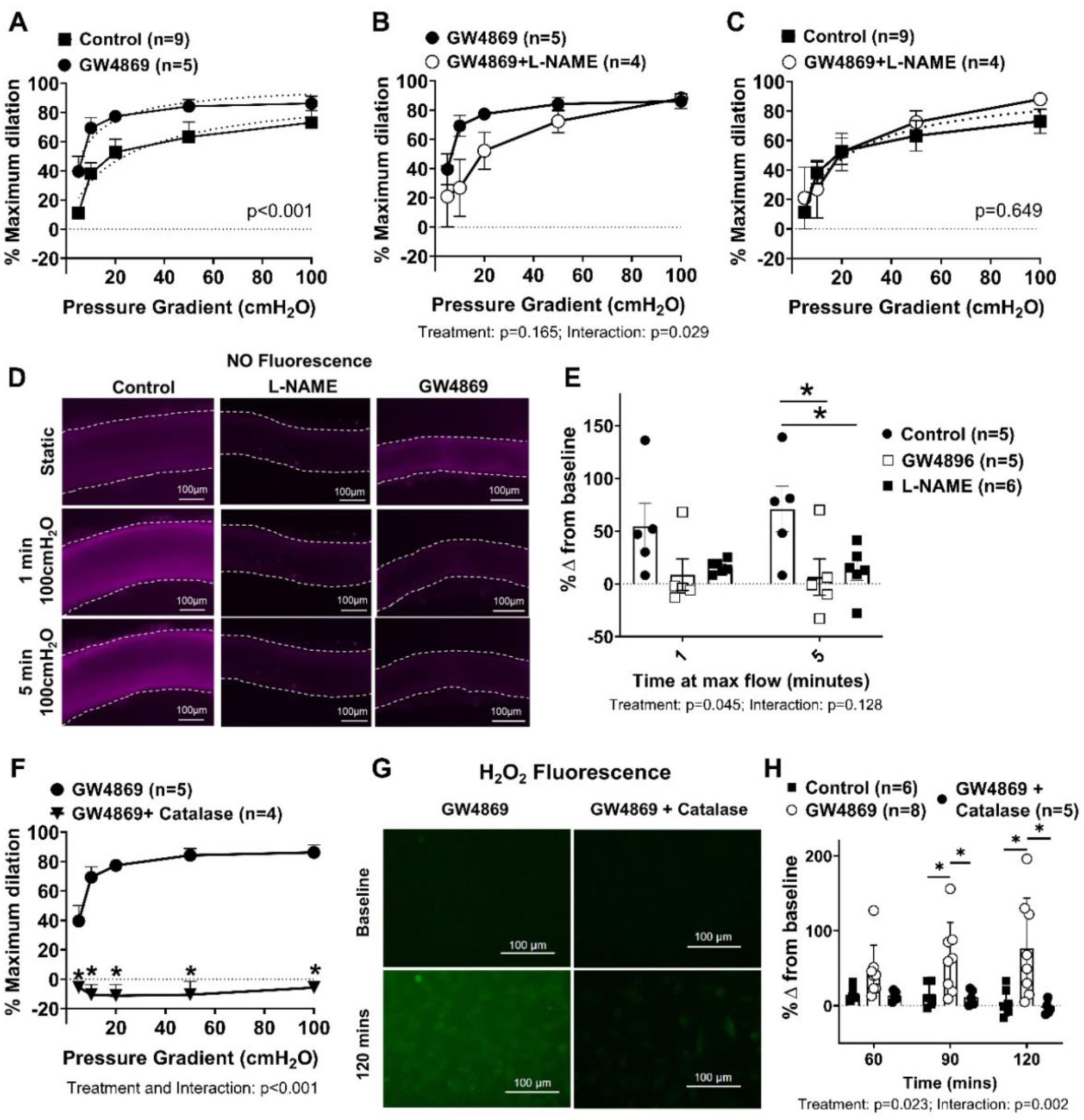
Inhibition of neutral sphingomyelinase impairs NO-mediated dilation to flow in arterioles from healthy (nonCAD) patients. (A) Response to flow in arterioles from nonCAD patients with (n=5) and without (n=9) NSmase inhibitor treatment (GW4869, 4μM, 30mins). (B) Response to flow in arterioles treated with GW4869 with (n=4) or without (n=5) L-NAME (100μM). (C) Flow response in GW4869+L-NAME (n=4) treated vessels compared to untreated control (n=9). (D) Representative images of untreated vessels (n=5) and vessels treated with L-NAME (n=6) or GW4869 (n=4; 30mins all), after intraluminal incubation with a nitric oxide fluorescence probe. Dotted white lines show outline of vessels. (E) Quantification of percent change in nitric oxide fluorescence intensity from baseline to 1 and 5 minutes of maximal flow (100cmH_2_O). (F) Flow response following treatment with catalase (500U/mL) and GW4869 (n=5) compared to GW4869 alone (n=4). (G) Representative images of HUVECs treated with GW4869 with and without catalase and exposed to a H_2_O_2_ (peroxy yellow 1) fluorescence probe. For visualization, a 40% increase in both the brightness and contrast was applied to the representative images. (H) Quantification of change in H_2_O_2_ fluorescence over time for up to 120 mins (control n=6, GW4869 n=8, GW4869+catalase n=5). (A,C) – non-linear regression with a 4-parameter fit (min constrained to 0, max constrained to 100, and EC50 constrained to >0); dotted line represents best-fit. (B, E, F, H) – two-way repeated measured ANOVA with Holm-Sidaak multiple comparisons test. *p<0.05

To confirm the loss of NO production in response to shear during NSmase inhibition, NO fluorescent imaging was performed in human arterioles exposed to maximal flow (pressure gradient of 100 cmH2O). As shown in Figure 1D-E, acute inhibition of NSmase (GW4869: 4μM, 30min) reduced the amount of NO generated in arterioles from nonCAD subjects exposed to maximal flow for 5 minutes (DBM control vs GW4869 at 5 mins, n=5 both: -64.40±22.18, p=0.015). Signal specificity was confirmed in arterioles treated with L-NAME (DBM control vs L-NAME at 5 mins -58.13±21.23; p=0.015, n=5 & 6, respectively). Flow-induced generation of NO was unaffected in arterioles exposed to vehicle alone (Supplemental Figure 1C).

To test whether inhibition of NSmase resulted in endothelial dysfunction, defined as a switch in the mediator of FID from NO to H_2_O_2_, vessels were treated acutely with the NSmase inhibitor GW4869 in the presence of catalase. This diminished dilation to flow (DBM GW4869, n=5 vs GW4869+catalase, n=4: -79.95±7.64, p<0.001; Figure 1F). To determine whether inhibition of ceramide formation directly contributes to the production of endothelial H_2_O_2_, HUVECs were treated with GW4869 (4μM) and the amount of H_2_O_2_ generated over time was measured using the fluorescent probe PY1. An increase in H_2_O_2_ fluorescence was observed over time up to 120 minutes in cells treated with GW4869 (DBM at 120mins; Control, n=6 vs GW4869, n=8: 71.64±19.07, p<0.001; Figure 1G-H), suggesting that loss of ceramide formation upregulates endothelial H_2_O_2_ production independent of flow. The increase in H_2_O_2_ was diminished by the addition of catalase, confirming signal specificity (DBM 120mins; GW4869, n=8 vs GW4869+catalase, n=5: -79.77±20.13, p<0.001; Figure 1G-H). Vehicle control had no effect on H_2_O_2_ production over time (Supplemental Figure 1D).

### Exogenous ceramide promotes acute NO production and restores endothelial function during NSmase inhibition in nonCAD arterioles

To determine whether ceramide can generate NO in human microvessels, arterioles were exposed to increasing doses of C2-ceramide (10^-9^ to10^-5^) in the presence of the NOS inhibitor L-NAME or NO-scavenger C-PTIO. As shown in figure 2A, exposure to increasing doses of the active C2-ceramide resulted in microvascular dilation as opposed to the biologically inactive dihydroceramide and vehicle control (DBM at 10^-5^; ceramide vs DMSO, n=5 both: 27.68±9.89, p=0.049; dihydroceramide, n=4 vs DMSO, n=5: -8.32±8.45, p=0.370). The presence of either L-NAME or C-PTIO significantly reduced the ability of the arterioles to dilate to C2-ceramide (DBM at 10^-5^; ceramide, n=5 vs ceramide+C-PTIO, n=4: -31.13±9.38, p=0.016; ceramide, n=5 vs ceramide+L-NAME, n=4: -33.40±10.06, p=0.017; Figure 2B).

**Figure 2.**
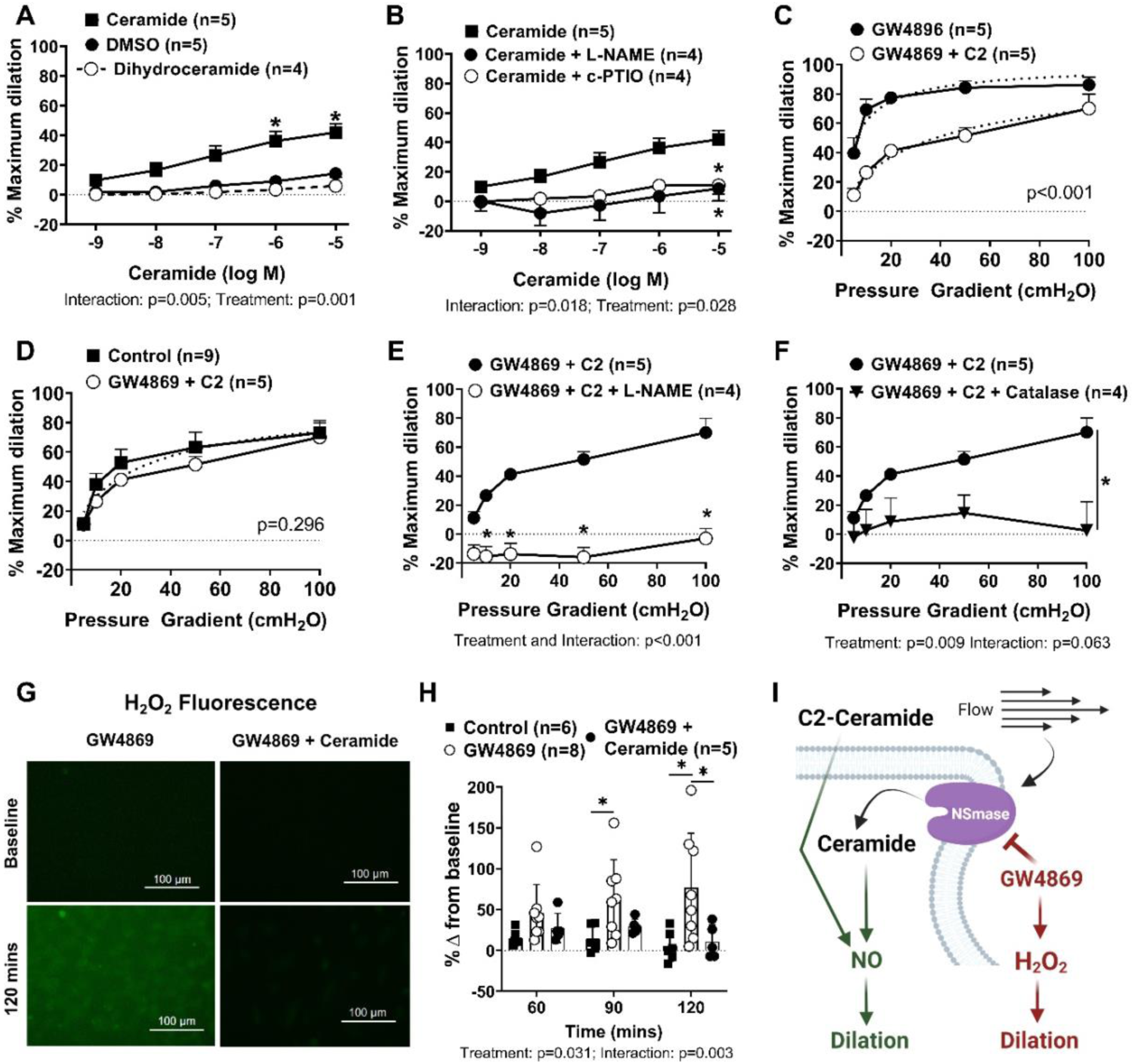
Exogenous ceramide promotes acute NO production and restores endothelial function during NSmase inhibition in healthy (nonCAD) patients. (A) Exogenous C2-ceramide (n=5) induced dilation (10^-9^ to 10^-5^ M; 1min) compared to vehicle (DMSO; n=5) and dihyro-C2-ceramide (n=4) in arterioles from healthy, nonCAD patients. (B) Ceramide-induced dilation with L-NAME (100μM; n=4) and CPTIO (100μM; n=4; 30min all) compared to ceramide alone (n=5). Flow response with exogenous ceramide (10μM, 30mins) + GW4869 (n=5) compared to (C) GW4869 alone (4μM, 30mins, n=5) and (D) untreated control vessels (n=9). Flow response in vessels treated with ceramide+GW4869 (n=4), in the presence of (E) L-NAME (n=4) and (F) catalase (500U/ml; n=4). (G) Representative images of HUVECs treated with GW4869 with and without C2-Ceramide and exposed to a H_2_O_2_ (peroxy yellow 1) fluorescence probe. For visualization, a 40% increase in both the brightness and contrast was applied to the representative images. (H) Quantification of change in H_2_O_2_ fluorescence over time for up to 120 mins (Control n=6, GW4869 n=8, GW4869+Ceramide n=5). (I) Schematic representation of the role of acute ceramide formation in maintaining NO-mediated dilation. Schematic was created using Biorender.com and permission was obtained for publication. (A,B,E,F,H) – two-way repeated measured ANOVA with Holm-Sidaak multiple comparisons test. (C,D) – non-linear regression with a with a 4-parameter fit (min constrained to 0, max constrained to 100, and EC50 constrained to >0); dotted line represents best-fit. *p<0.05

To answer the question of whether exogenous ceramide can restore endothelial function to arterioles unable to produce ceramide via NSmase, C2-ceramide (10μM, 30mins) was administered simultaneously with GW4869 which resulted in a vasodilatory flow pattern similar to control vessels (Figure 2C-D; 95% CI EC50 of best fit curve for control, n=9 and GW4869+ceramide, n=5: 21.18-38.51, p=0.296). NO-mediated FID was also restored in arterioles exposed to both GW4869 and C2-ceramide as dilation to flow was significantly reduced with the addition of L-NAME (DBM GW4869+ceramide, n=5 vs GW4869+ceramide+L-NAME, n=4: -52.48±5.91, p<0.001; Figure 2E). Surprisingly, FID was also reduced in the presence of catalase (DBM GW4869+ceramide, n=5 vs GW4869+ceramide+catalase, n=4: -34.86±9.83, p=0.009; Figure 2F). In line with these findings, acute ceramide induced dilation was inhibited in the presence of catalase (Supplemental Figure 1E), suggesting that C2-ceramide-induced NO production may require formation of H_2_O_2_.

As shown in figure 2G-H, overall H_2_O_2_ production was diminished in HUVECs treated with both GW4869 and ceramide (DBM at 120mins; GW4869, n=8 vs GW4869+ Ceramide, n=5: -66.16±20.53, p=0.002). The role of acute ceramide formation in maintaining NO-mediated dilation is schematized in figure 2I.

### The beneficial role of ceramide on nonCAD microvascular endothelial function is dependent on formation of S1P

To test whether the beneficial effects of ceramide formation on endothelial function are due to its metabolite sphingosine-1-phosphate (S1P), changes in vessel diameter were measured in response to C2-ceramide during simultaneous inhibition of sphingosine kinase (sphingosine kinase inhibitor; SpK-I). The vasodilatory response to exogenous C2-ceramide was reduced in arterioles pre-treated with SpK-I (1μM, 30min) as shown in Figure 3A (DBM at 10^-5^ M; ceramide, n=5 vs ceramide+SpK-I, n=6: -29.27±7.25, p=0.021).

**Figure 3.**
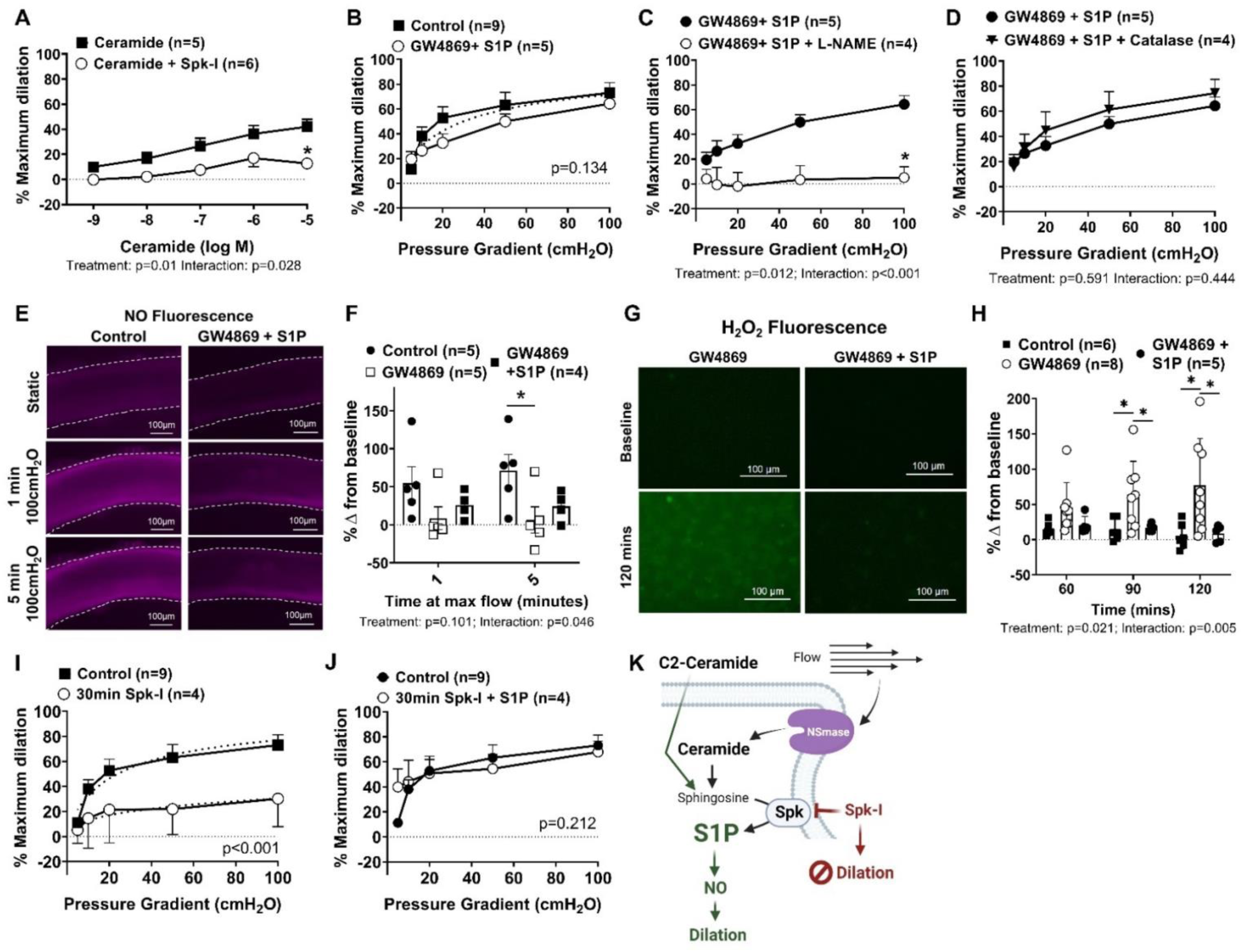
The beneficial role of ceramide on healthy (nonCAD) microvascular endothelial function is dependent on formation of S1P. (A) C2-Ceramide-induced dilation (10^-9^ to 10^-5^ M, 1min) in vessels treated with sphingosine-kinase inhibitor (SpK-I 1μM, 30mins; n=4) versus control (n=5). (B) Flow response in vessel treated with exogenous S1P (1μM, 30mins) + GW4869 (4μM, 30mins; n=5) compared to untreated control (n=9). Response to flow in vessels treated with S1P+GW4869 (n=5), in the presence of (C) L-NAME (100μM; n=4) and (D) catalase (500U/ml; n=4). (E) Representative images of untreated vessels (n=5) and vessels treated with GW4869+S1P (30mins all; n=4), after intraluminal incubation with a nitric oxide fluorescence probe. Dotted white lines show outline of vessels. (F) Quantification of percent change in nitric oxide fluorescence from baseline to 1 and 5 minutes of maximal flow (100cmH_2_O). (G) Representative images of HUVECs treated with GW4869 with and without S1P and exposed to a H_2_O_2_ (peroxy yellow 1) fluorescence probe. For visualization, a 40% increase in both the brightness and contrast was applied to the representative images. (H) Change in H_2_O_2_ fluorescence intensity for up to 120 mins (control n=6, GW4869 n=8, GW4869+S1P n=5). (I) Response to flow with inhibition of sphingosine-kinase (30mins; n=4) compared to control (n=9) (J) Flow response with SpK-I+S1P (n=4; 30min) compared to control (n=9). (K) Schematic representation of the role of S1P formation in ceramide-induced NO-signaling. Schematic was created using Biorender.com and permission was obtained for publication. (A, C, D, F, H) – two-way repeated measured ANOVA with Holm-Sidaak multiple comparisons test. (B, I, J) – non-linear regression with a 4-parameter fit (min constrained to 0, max constrained to 100, and EC50 constrained to >0); dotted line represents best-fit. *p<0.05

Similar to treatment with C2-ceramide, acute exposure to S1P restored the vasodilatory flow pattern in vessels during inhibition of NSmase to that of what is observed in untreated nonCAD arterioles (Figure 3B, 95% CI EC50 of best fit curve for control, n=9 and GW4869+S1P, n=5: 21.92-45.54, p=0.134). S1P also restored NO-mediated FID as dilation was impaired in the presence of L-NAME (DBM GW4869+S1P, n=5 vs GW4869+S1P+L-NAME, n=4: -36.62±10.89, p=0.021; Figure 3C). A striking difference between administration of C2-ceramide versus S1P during NSmase inhibition is that treatment with S1P restored NO as the primary mediator of FID as dilation to flow was maintained in the presence of catalase (DBM GW4869+S1P, n=5 vs GW4869+S1P+catalase, n=4: -6.61±11.74, p=0.591;Figure 3D).

Production of NO in vessels treated with GW4869 and S1P was confirmed by measuring changes in NO fluorescence in response to flow, which showed no difference compared to control vessels (Figure 3E-F; DBM control, n=5 vs GW4869+S1P, n=4 at 5 min: 46.55±25.28, p=0.079). As shown in Figure 3G-H, the upregulation in H_2_O_2_ production with NSmase inhibition over time was diminished when cells were treated with GW4869 and S1P (DBM at 120mins; GW4869, n=8 vs GW4869+ S1P, n=5: -68.57±20.19, p=0.001).

The impact of inhibiting SpK and reducing intracellular production of S1P on FID was then examined by acutely administering SpK-I (1μM, 30min) prior to flow. Interestingly, rather than initiating a switch from NO-to H_2_O_2_-mediated FID, there was a significant reduction in overall dilation to flow (95% CI EC50 of best fit curves for control, n=9: 15.74-37.34 vs SpK-I, n=4: 56.66-undefined, p<0.001; Figure 3I), which was restored with the administration of exogenous S1P (Figure 3J; 95% CI EC50 of best fit curves for Control, n=9 and SpK-i+S1P, n=4 15.32-35.74; p=0.212). FID was unaffected when exposed to vehicle control (supplemental Figure 1A). The role of S1P in promoting NO via ceramide is schematized in Figure 3K.

### Inhibition of S1PR1 impairs NO-mediated dilation to flow in nonCAD arterioles

As shown in Figure 4A, ceramide-induced dilation is impaired when treated with W146 (10μM, 30mins), an inhibitor of S1PR1 (DBM at 10^-5^ M; ceramide, n=5 vs ceramide+W146, n=4 -39.73±5.99, p=0.010).

**Figure 4.**
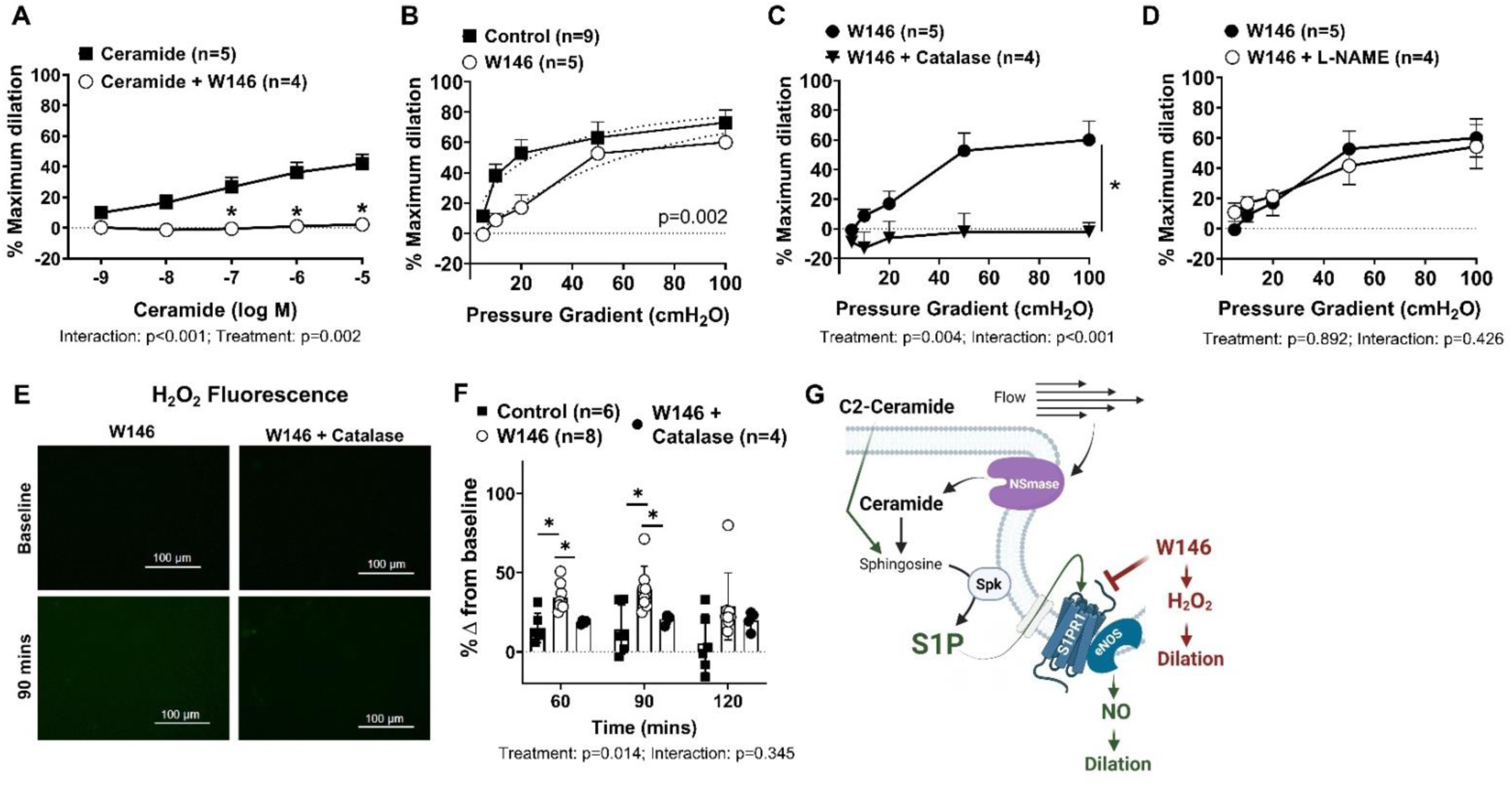
Inhibition of S1PR1 impairs NO-mediated dilation to flow in healthy (nonCAD) arterioles. (A) C2-Ceramide-induced dilation (10^-9^ to 10^-5^ M, 1min) in vessels treated with a S1P-receptor-1 inhibitor (W146 10μM, 30mins; n=4) compared to control (n=5). (B) Flow response following W146 treatment (n=5; 30mins) compared to untreated control (n=9). Response to flow in vessels treated with W146 (n=5), in the presence of (C) catalase (500U/ml; n=4) and (D) L-NAME (100μM; n=4). (E) Representative images of HUVECs treated with W146 with and without catalase and exposed to a H_2_O_2_ (peroxy yellow 1) fluorescence probe. For visualization, a 40% increase in both the brightness and contrast was applied to the representative images. (F) Quantification of H_2_O_2_ fluorescence intensity over time for up to 120 mins (control n=6, W146 n=8, W146+catalase n=4). (G) Schematic representation of the role of S1PR1 in promoting NO production. Schematic was created using Biorender.com and permission was obtained for publication. (A, C, D, F) – two-way repeated measured ANOVA with Holm-Sidaak multiple comparisons test. (B) non-linear regression with a 4-parameter fit (min constrained to 0, max constrained to 100, and EC50 constrained to >0); dotted line represents best-fit. *p<0.05

To investigate the role of S1PR1 in FID, arterioles were also treated acutely with W146 (10μM, 30mins) prior to the initiation of flow. Inhibition of S1PR1 delayed dilation to flow (95% CI EC50 of best fit curves for control, n=9: 15.74-37.34 vs W146, n=5: 41.18-98.71; Figure 4B), an effect not observed in vessels treated with vehicle control (Supplemental Figure 2B). In microvessels treated with the S1PR1 inhibitor, response to flow was impaired during exposure to catalase (Figure 4C; DBM W146, n=5 vs W146+catalase, n=4: -34.0±7.99; p=0.004) but not those treated with L-NAME (Figure 4D; DBM W146, n=5 vs W146+L-NAME, n=4: - 1.39±9.95, p=0.892). NO-mediated FID was maintained in vessels treated with vehicle control alone (Supplemental Figure 2C). In cultured HUVECs, inhibition of S1PR1 increased H_2_O_2_ at 60 and 90 min (DBM at 90mins; control vs W146 25.29±8.13, p=0.015; Figure 4E), which was not sustained beyond 90 min. This increase in H_2_O_2_ was diminished with the addition of catalase (DBM at 90mins; W146, n=8 vs W146 +catalase, n=4: -18.83±5.34, p=0.015; Figure 4F). The role of S1PR1 signaling in promoting NO is schematized in Figure 4G.

### Agonism of S1PR1 restores NO-dependent dilation during NSmase inhibition in arterioles from nonCAD patients

The vasodilatory pattern to flow that is observed in healthy, nonCAD control vessels was restored in nonCAD arterioles treated with CYM5442, an agonist of S1PR1, during NSmase inhibition (Figure 5A, 95% CI EC50 of best fit curves for Control, n=9 and GW4869+CYM5442, n=4: 14.02-30.30, p=0.175; supplemental Figure 2D). Agonism of S1PR1 in the presence of NSmase inhibitor also restored dependency on NO for FID as L-NAME diminished dilation (Figure 5B; DBM GW4869+CYM5442, n=4 vs GW4869+CYM5442+L-NAME, n=4: -70.75±19.88, p=0.012), however dilation was also reduced with catalase (Figure 5C; DBM GW4869+CYM5442, n=4 vs GW4869+CYM5442+catalase, n=4: -52.05±11.69, p=0.004). Figure 5D-E show that endothelial H_2_O_2_ production during NSmase inhibition may be diminished when treated with CYM5442 (DBM at 120mins; GW4869, n=8 vs GW4869+CYM5442, n=5; -49.90±23.86, p=0.07). The role of S1PR1 in promoting NO downstream of ceramide formation is schematized in figure 5F.

**Figure 5.**
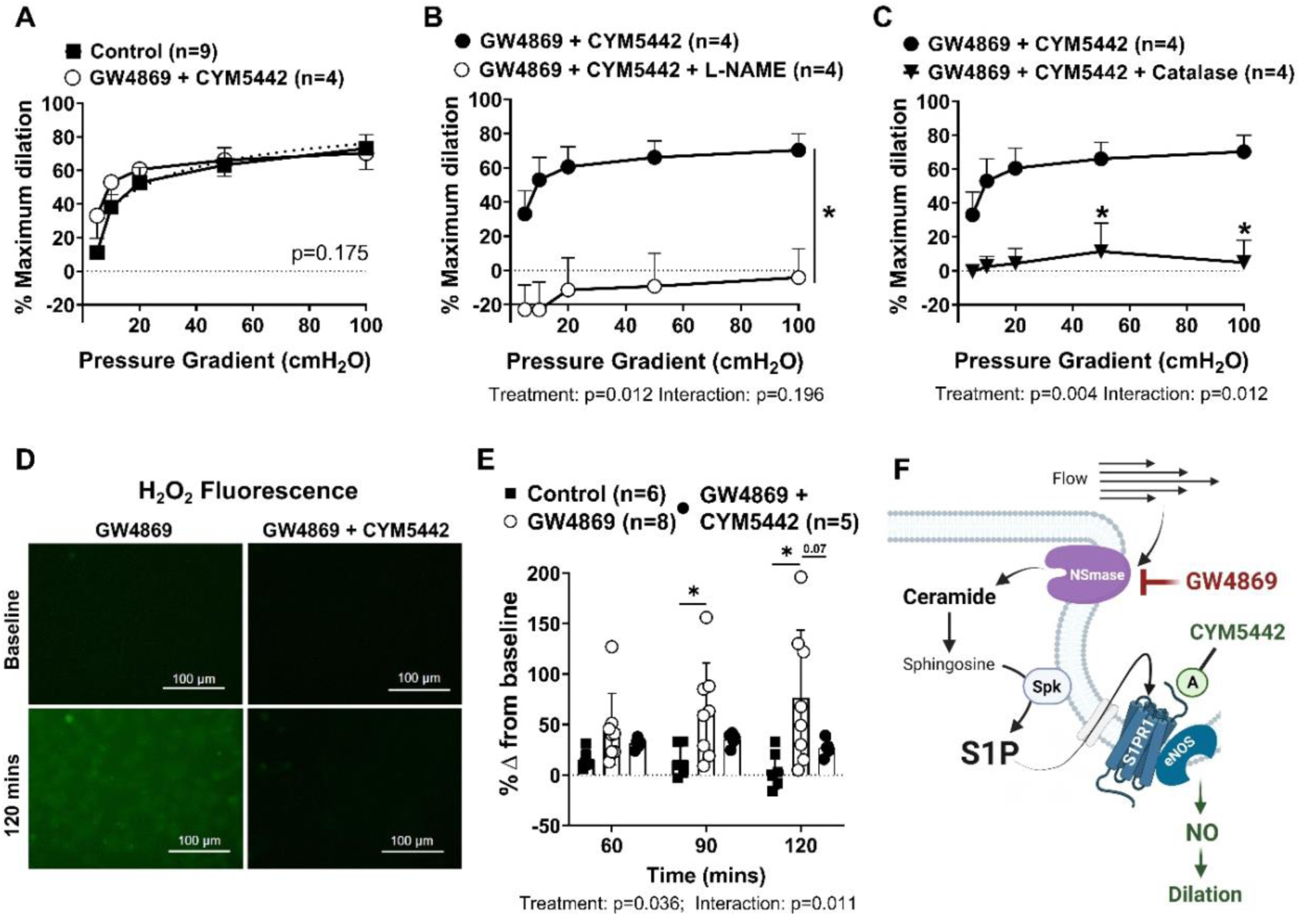
Agonism of S1PR1 restores NO-dependent dilation during NSmase inhibition in arterioles from nonCAD patients. (A) Flow response in vessels treated with an agonist of S1PR1 (CYM5442, 1μM, 30mins) + GW4869 (4μM, 30mins; n=4) compared to untreated control (n=9). Response to flow in vessels treated with CYM5442+GW4869 (n=4), in the presence of (B) L-NAME (100μM; n=4) and (C) catalase (500U/ml; n=4). (D) Representative images of HUVECs treated with GW4869 with and without CYM5442 and exposed to a H_2_O_2_ (peroxy yellow 1) fluorescence probe. For visualization, a 40% increase in both the brightness and contrast was applied to the representative images. (E) Quantification of change in H_2_O_2_ fluorescence over time for up to 120 mins (control n=6, GW4869 n=8, GW4869+CYM5442 n=5). (F) Schematic representation of the role of S1PR1 in promoting NO downstream of ceramide formation. Schematic was created using Biorender.com and permission was obtained for publication. (A) – non-linear regression with a 4-parameter fit (min constrained to 0, max constrained to 100, and EC50 constrained to >0); dotted line represents best-fit. (B, C, E) – two-way repeated measured ANOVA with Holm-Sidaak multiple comparisons test. *p<0.05

### S1PR1 and S1PR3 receptors have compensatory signaling during flow and inhibition of S1PR3 results in a dual dependency on both NO and H_2_O_2_ for dilation

We assessed whether the dual dependency on both NO and H_2_O_2_ during S1PR1 agonism is reflective of normal S1PR1 signaling in human nonCAD arterioles. S1PR1 and S1P-receptor-3 (S1PR3) receptors have compensatory signaling during flow in nonCAD vessels as maximal dilation is preserved during inhibition of S1PR3 (CAY10444: 10μM, 30mins) (Figure 6A, 95% CI EC50 of best fit curves for Control, n=9 and CAY10444, n=4: 18.72-34.60, p=0.520) or S1PR1 (Figure 4B) alone, while the dual inhibition of S1PR1 and S1PR3 diminished overall vasodilation to flow (Figure 6B, 95% CI EC50 of best fit curves for Control, n=9: 15.74-37.34 vs W146+CAY10444, n=5: 95.65-69794, p<0.001). Inhibition of S1PR3, which presumably prioritizes endothelial signaling through S1PR1, resulted in a dual dependency on both NO and H_2_O_2_ for FID as both L-NAME (Figure 6C; DBM CAY10444, n=4 vs CAY10444+L-NAME, n=5: -40.93±14.58, p=0.027) and catalase (Figure 6D; DBM CAY10444, n=4 vs CAY10444+catalase, n=5: -24.23±9.47, p=0.038) reduced dilation to flow.

**Figure 6.**
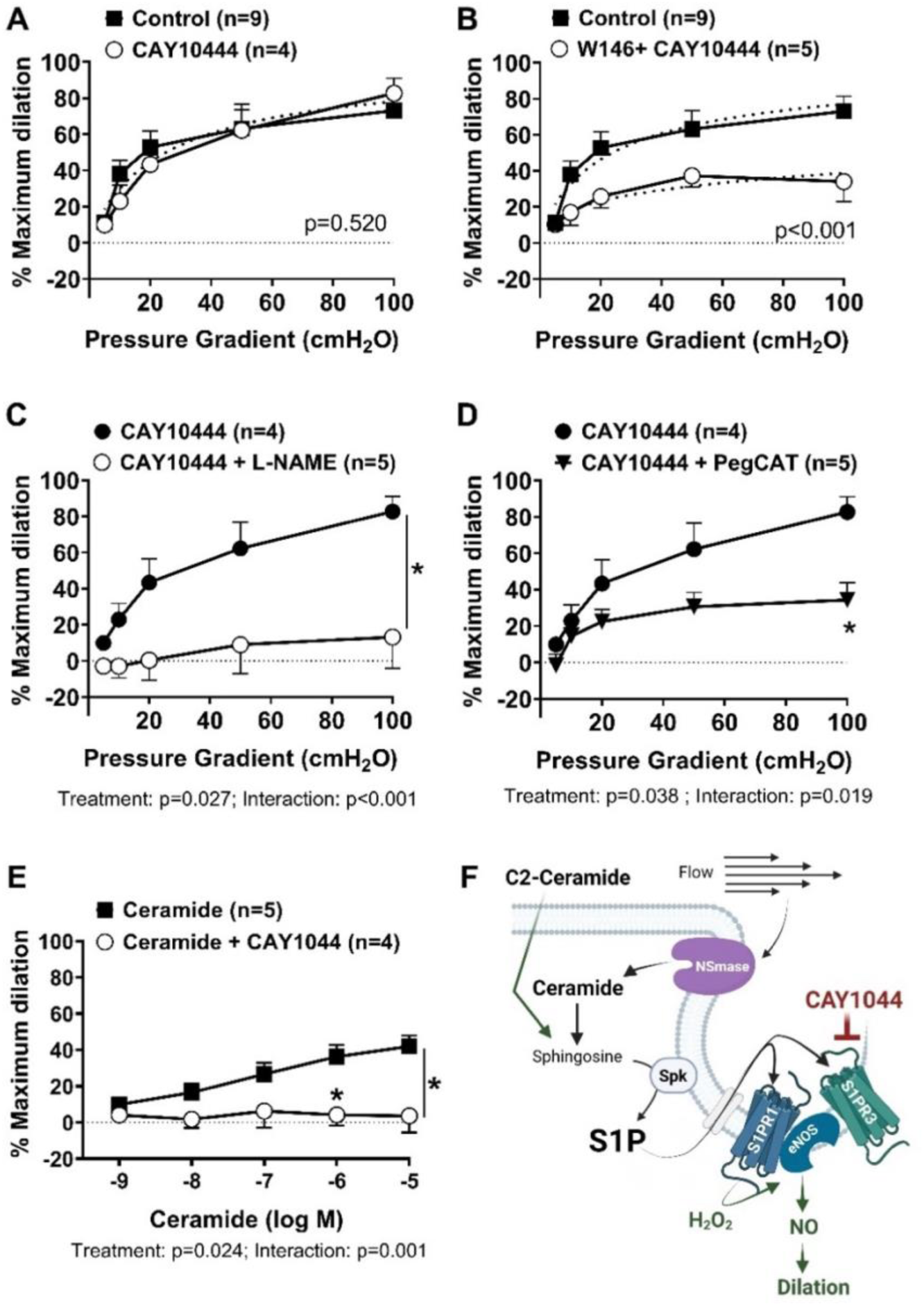
S1PR1 and S1PR3 receptors have compensatory signaling during flow. Inhibition of S1PR3 results in a dual dependency on both NO and H_2_O_2_ for dilation. Response to flow in arterioles from nonCAD patients treated with (A) S1PR1 inhibitor CAY10444 (10μM; n=4; 30min), and (B) CAY10444+S1PR1-inhibitor W146 (10μM; n=5; 30min) compared to control (n=9). Response to flow in vessels treated with CAY10444 (n=4), in the presence of (C) L-NAME (100μM; n=5) and (D) catalase (500U/ml; n=5). (E) C2-Ceramide-induced dilation (10^-9^ to 10^-5^ M, 1min) in vessels treated with CAY10444 (n=4) compared to control (n=5). (F) Schematic Representation of the role of S1PR1 and S1PR3 in ceramide signaling in nonCAD arterioles. Schematic was created using Biorender.com and permission was obtained for publication. (A, B) – non-linear regression with a 4-parameter fit (min constrained to 0, max constrained to 100, and EC50 constrained to >0); dotted line represents best-fit. (C, D, E) – two-way repeated measured ANOVA with Holm-Sidaak multiple comparisons test. *p<0.05

To assess if S1PR3 plays a role in ceramide-induced NO production, nonCAD arterioles were treated with CAY10444 (10μM, 30mins) and exposed to increasing doses of C2-ceramide. Inhibition of S1PR3 impaired ceramide-induced dilation (Average DBM 10^-9^ - 10^-5^ M; ceramide, n=5 vs ceramide+CAY10444, n=4: -22.25±7.72, p=0.001). The role of S1PR1 and S1PR3 in ceramide signaling in nonCAD arterioles is schematized in figure 6F.

### Acute ceramide induced dilation in CAD arterioles is dependent on H_2_O_2_ production through S1P/S1PR3 signaling

In order to examine the ceramide/S1P signaling axis in the human microvasculature during disease (CAD), isolated arterioles from patients formally diagnosed with CAD were subjected to increasing doses of C2-ceramide. Dilation to ceramide was similar in vessels from nonCAD and CAD patients (Supplemental Figure 5A). Unlike in vessels from healthy patients, the response to C2-ceramide was preserved in vessels from CAD patients when exposed to either L-NAME or CPTIO (DBM at 10^-5^ M; ceramide, n=4 vs ceramide+C-PTIO, n=5: -18.50±13.45, p=0.337; ceramide, n=5 vs ceramide+L-NAME, n=7: -10.91±11.96, p=0.573, Figure 7A).

**Figure 7.**
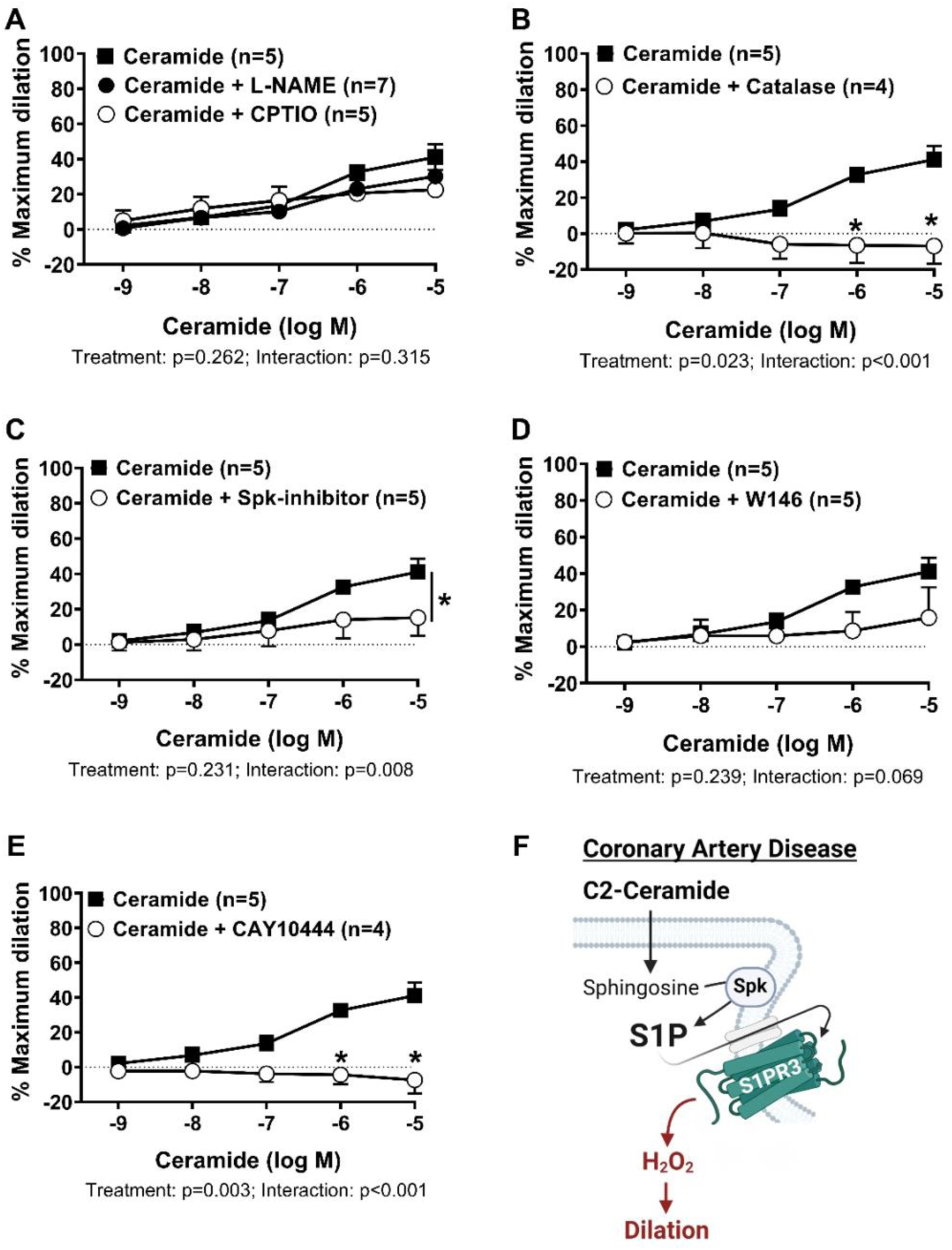
Acute ceramide induced dilation in CAD arterioles is dependent on H_2_O_2_ production through S1P/S1PR3 signaling. Exogenous C2-ceramide (n=5) induced dilation (10^-9^ to 10^-5^ M; 1min) of arterioles from patients with CAD in the presence of (A) L-NAME (100μM; n=7), CPTIO (100μM; n=5; 30min all), or (B) Catalase (500U/ml). Ceramide-induced dilation of vessels from patients with CAD treated with (C) sphingosine-kinase inhibitor (SpK-I 1μM, 30mins; n=5), (D) S1PR1 inhibitor (W146 10μM, 30mins; n=5), and (E) S1PR3 inhibitor (CAY10444 10μM, 30mins; n=4). (F) Schematic representation of the shift towards S1PR3-mediated signaling in response to ceramide in arterioles from patients with CAD. Schematic was created using Biorender.com and permission was obtained for publication. (A-E) – two-way repeated measured ANOVA with Holm-Sidaak multiple comparisons test. *p<0.05

However, ceramide-induced dilation was reduced during treatment with catalase (DBM at 10^-5^ M; ceramide, n=5 vs ceramide+ catalase, n=4: -48.03±9.33, p<0.001;Figure 7B). Vasodilation to increasing doses of ceramide was also reduced during inhibition of sphingosine kinase (Average DBM 10^-9^ - 10^-5^ M; ceramide, n=5 vs ceramide+ SpK-I, n=5: -11.00±8.50, p=0.008, Figure 7C), however was maintained during blockade of S1PR1 (DBM at 10^-5^ M; ceramide, n=5 vs ceramide+ W146, n=5 -25.18±18.22, p=0.711; Figure 7D). Rather, the response to ceramide was impaired during inhibition of S1PR3 (DBM at 10^-5^ M; ceramide, n=5 vs ceramide+ CAY10444, n=4: -48.45±10.73, p=0.015, Figure 7E), suggesting a shift towards S1PR3-mediated signaling in response to ceramide in arterioles from patients with CAD (schematized in Figure 7F).

### Ceramide formation remains a critical step in FID during disease

To examine the importance of ceramide formation during flow in the human microvasculature during disease (CAD), isolated arterioles from patients formally diagnosed with CAD were exposed to increasing flow rates during NSmase inhibition. Treatment with GW4869 (4μM, 30min) diminished dilation to flow (Figure 8A; 95% CI EC50 of best fit curve for control, n=5, 17.02-29.82 vs GW4869, n=5, 63.81-2.7E9; p<0.001). Treatment with exogenous ceramide during NSmase inhibition increased dilation to lower flow rates but did not fully restore FID (Figure 8B, 95% CI EC50 of best fit curve for control, n=5, 17.02-29.82 vs GW4869+C2, n=5, 11.39 - undefined; p<0.001; supplemental Figure 5B). FID remained diminished when treated with exogenous S1P (Figure 8C; 95% CI EC50 of best fit curve for control, n=5, 17.02-29.82 vs GW4869+S1P, n=4, 90.48 - undefined; p<0.001; supplemental Figure 5C) during NSmase inhibition, suggesting that in diseased vessels the ceramides during flow may primarily signal through pathways alternative to S1P.

**Figure 8.**
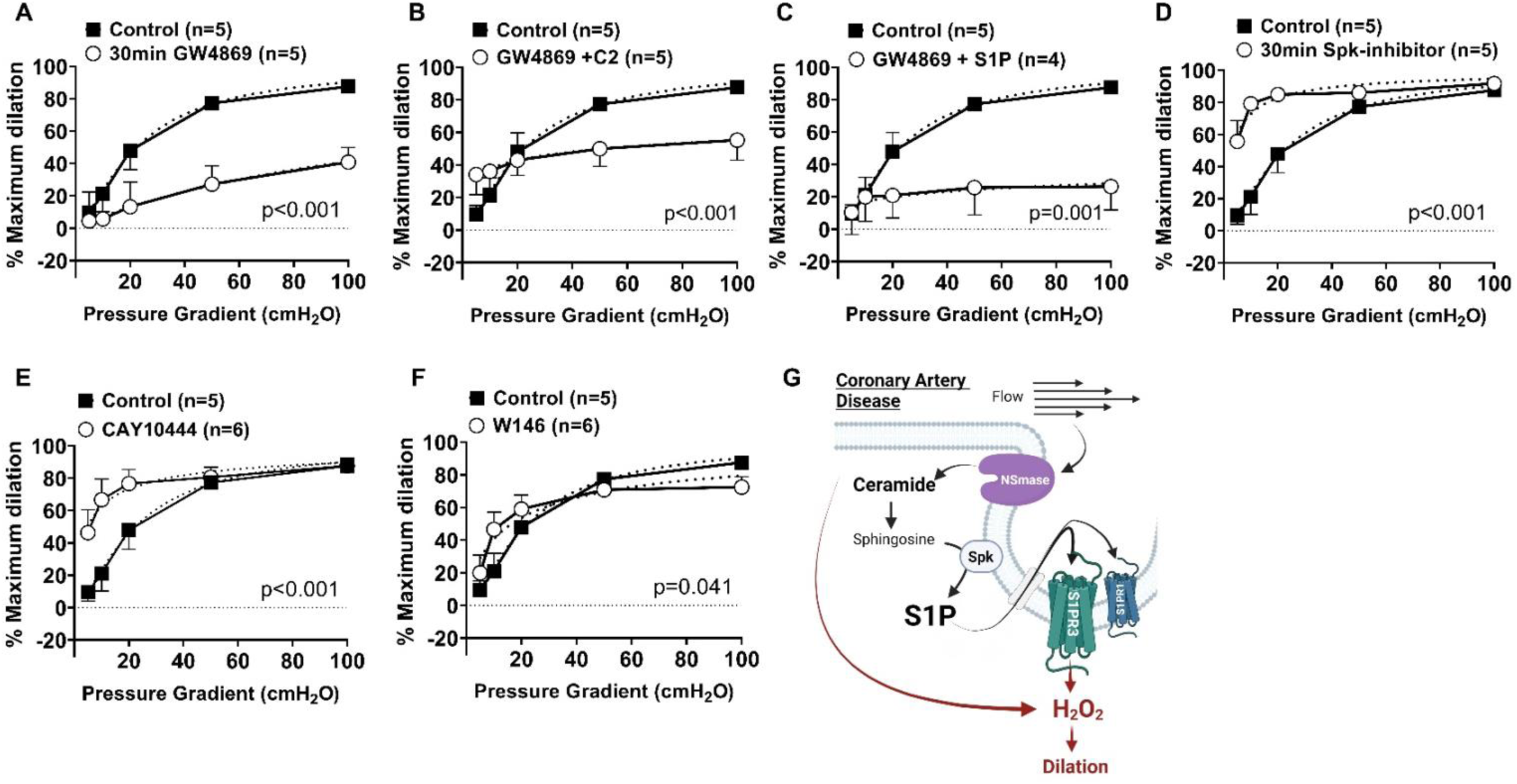
Ceramide formation remains a critical step in FID during disease. (A) Response to flow in arterioles from CAD patients treated with a NSmase-inhibitor (GW4869, 4μM; n=5) compared to control (n=5). (B) Flow response with exogenous ceramide (10μM, 30mins) + GW4869 (n=5) compared to control (n=5). (C) Flow response with exogenous S1P (1μM, 30mins) + GW4869 (n=4) compared to control (n=5). Response to flow in arterioles from patients with CAD treated with (D) sphingosine-kinase inhibitor (SpK-I, 1μM, 30mins, n=5), (E) S1PR3 inhibitor (CAY10444, 10μM, 30mins, n=6), and (F) S1PR1 inhibitor (W146, 10μM, 30mins, n=6) compared to control (n=5). (G) Schematic showing the role of ceramide and S1P during flow in diseased vessels. Schematic was created using Biorender.com and permission was obtained for publication. (A-F) – non-linear regression with a 4-parameter fit (min constrained to 0, max constrained to 100, and EC50 constrained to >0); dotted line represents best-fit. *p<0.05

To confirm that S1P formation does not play a crucial role in flow-signaling in disease arterioles, vessles were treated with SpK-I (1μM, 30mins) and exposed to increasing flow rates. Although maximal dilation to flow was maintained during sphingosine kinase inhibition, treatment with SpK-I heightened vessel dilation at lower flow rates (Figure 8D, 95% CI EC50 of best fit curve for control, n=5, 17.02-29.82 vs SpK-I, n=5, 0.48 - 5.37; p<0.001). A similar response was also seen with S1PR3 inhibition (Figure 8E, 95% CI EC50 of best fit curve for control, n=5, 17.02-29.82 vs CAY10444, n=6, 0.42 - 9.38; p<0.001), suggesting that S1P/S1PR3 signaling may play a role in mediating flow-response during lower flow rates in diseased vessels. Inhibition of S1PR1 altered response to flow, however to a lesser degree (Figure 8F, 95% CI EC50 of best fit curve for control, n=5, 17.02-29.82 vs W146, n=6, 9.47 – 27.51; p<0.001). figure 8G schematizes the role of ceramide and S1P during flow in diseased vessels.

## DISCUSSION

Microvascular endothelial dysfunction, both in the peripheral and coronary circulation, independently predict major adverse cardiovascular events (MACE)^26–28^ and can precede the development of large vessel disease^27, 29^. Understanding mechanisms contributing to microvascular homeostasis versus dysfunction is crucial for developing novel therapeutic strategies for reducing cardiovascular morbidity and mortality^30^. The major findings of this study are 6-fold:1) Acute Nsmase-mediated ceramide formation is necessary for maintaining NO-mediated FID in arterioles from otherwise healthy adults; 2) Endothelial cells produce H_2_O_2_ during static conditions when NSmase is inhibited; 3) Ceramide conversion to S1P and activation of S1PR1/3 is required for ceramide-induced NO-production; 4) NSmase-mediated ceramide formation is not only necessary to produce NO in response to flow, but for maintaining an appropriate response to flow at lower flow rates; 5) In arterioles from patients with CAD, inhibition of NSmase-mediated ceramide formation impairs dilation to flow; 6) Arterioles from patients with CAD primarily signal through S1PR3 in response to ceramide, resulting in dilation dependent on generation of H_2_O_2_ as opposed to NO. The data presented here suggests that despite key differences in downstream signaling between health and disease, acute NSmase-mediated ceramide formation and its subsequent conversion to S1P is necessary for proper functioning of the human microvascular endothelium.

The endothelium relies heavily on sphingolipid-rich invaginated microdomains to both sense changes in blood flow and transduce external signals by stimulating formation of vasodilator compounds^31, 32^. Czarny et al. have previously shown that NSmase acts as a mechanosensor within caveolae and is capable of rapidly increasing endothelial ceramide levels in a pressure- and flow rate-dependent manner^18, 33^. On the contrary, treatment with GW4869 decreases NSmase activity and results in accumulation of sphingomyelin and reduction in ceramides^34^ within the plasma membrane. Preclinical studies have indicated that ceramide is capable of activating eNOS^35, 36^ while inhibition reduces mechanotransduction of critical shear-sensing cascades^37–39^. Aside from acute ceramide formation through NSmase, other studies have highlighted the importance of chronic ceramide formation in generating NO via the *de novo* pathway, as inhibition of serine-palmitoyl transferase (rate-limiting enzyme in *de novo* pathway) reduces both dilation to flow as well as agonist-induced NO production in mice^40^. It is important to note here that reductions in NO signaling in animals often results in a decrease in vasodilatory capacity as opposed to the switch to H_2_O_2_ - mediated FID that is observed in humans. This compensatory pathway likely exists in humans to maintain critical flow regulation even in times of disease. Therefore, CVD preventative strategies that aim to significantly impair ceramide formation^16, 17, 41–43^ may actually prove detrimental to the human microvasculature.

Interestingly, acute inhibition of NSmase resulted in both H_2_O_2_-mediated FID as well as more rapid dilation at lower flow rates in arterioles from nonCAD patients. It is possible that suppressed ceramide formation, and/or accumulation of sphingomyelin results in unfavorable lipid composition and biophysical properties within the plasma membrane ultimately affecting the cell’s ability to sense and transduce flow changes^31^. It is unlikely that vessels treated with GW4869 were damaged as all arterioles maintained constriction following pre-constriction with ET-1 and dilated to over 70% of their maximal diameter in response to the smooth muscle dilator papaverine. Adding complexity to this observation is the fact that adding the NOS inhibitor L-NAME to the organ bath restored the normal flow response at lower pressure gradients. This suggests that the increased sensitivity to flow may be due to an exaggerated reactive oxygen species (ROS) response, for instance from uncoupled eNOS^44^. The fact that inhibition of NSmase in endothelial cells under static conditions increases H_2_O_2_ production over time supports this speculation as the H_2_O_2_ could contribute to eNOS uncoupling. Supplementation with ceramide, S1P, or an agonist of S1PR1 prevented the increase in basal endothelial H_2_O_2_ production during NSmase inhibition and restored the microvascular flow response at lower pressure gradients to what is observed in control vessels. This further supports that changes in ceramide signaling, rather than sphingomyelin accumulation/membrane biophysical properties, contribute to the exaggerated response to lower flowrates. In line with these findings, inhibition of S1P/S1PR1 signaling also impaired FID and significantly increased H_2_O_2_ production, albeit the increase in ROS was observed only up to 90 min as opposed to 120 min in cells treated with GW4869. This is not surprising as S1P/S1PR1 activation is downstream of ceramide formation via NSmase. Key questions remain as to exactly how the loss of ceramide/S1P/S1PR1 signaling stimulates production of endothelial ROS, even during the absence of flow.

The current study also sheds light on key differences in S1P signaling during flow between humans and preclinical models. Among the 5 major S1P receptors, and S1PR 1-3 are the most expressed within the vasculature with S1PR1 primary expressed in the vascular endothelium^25, 45^. S1PR2 is understood to be predominantly expressed in vascular smooth muscle cells whereas S1PR3 has been identified in both the endothelium and medial layer^25, 46^. We have previously shown that both S1PR1 and S1PR3 are indeed expressed in human microvessels from patients, both with and without CAD^25^. Cantalupo and Zhang et al. elegantly showed that following treatment with an inhibitor of S1PR1 (100nM and 1μM, 45 min), mouse mesenteric arteries exhibit decreased vasodilatory capacity during flow^47^. Here, we show that inhibition of S1PR1 acutely induces a switch to H_2_O_2_-mediated dilation. Again, the current results highlight the inter-species variation with regard to compensatory mechanisms of dilation exhibited in humans during decreased bioavailability of NO. This is further supported by our data showing that S1PR1 and S1PR3 have compensatory signaling during flow. Preclinical studies have linked S1PR1 activation with formation of NO^21^. However, here we show that agonism of S1PR1 during NSmase inhibition creates a dual dependency on both NO and H_2_O_2_ for FID of human arterioles. Similarly, inhibition of S1PR3, which presumably prioritizes signaling through S1PR1, also creates a dual dependency on NO and H_2_O_2_ production for FID. Together, these data suggest that H_2_O_2_ and NO are both produced downstream of S1PR1 signaling in healthy human arterioles. This is in agreement with our recent work showing that the mechanism of acute S1P-induced dilation in arterioles from nonCAD patients is dependent on production of both H_2_O_2_ and NO^25^.

Results show that arterioles from patients with CAD primarily signal through S1PR3 in response to ceramide, resulting in dilation dependent on generation of H_2_O_2_ as opposed to NO. Differences in G-protein coupled receptor (GPCR) signaling may account for the specific vasoactive mediator generated. Gi signaling associated with S1PR1 is known to favor NO formation whereas G12/13, and Gq, two of the three G proteins within S1PR3, are known to increase ROS^48–50^. Thus, a shift to signaling predominantly through S1PR3 may initiate or exaggerate endothelial dysfunction within the microvascular endothelium and increase the risk for future disease. Also unique to vessels from patients diagnosed with CAD, acute inhibition of ceramide formation via NSmase completely diminishes dilation to flow. This effect is not restored by addition of exogenous S1P. These arterioles also maintain vasodilatory capacity during suppression of the S1P/S1PR3 signaling axis, albeit with heightened dilation at lower flow rates. This suggests that during disease, dilation relies more on ceramide formation itself as opposed to its conversion to S1P and activation of S1PR3. Neutral ceramidase, an enzyme responsible for hydrolyzing ceramide to sphingosine, the backbone of S1P, is reduced in arterioles from patients with CAD^51^ which may be responsible for the accumulation of intracellular ceramide.

Growing evidence supports the hypothesis that chain length variations determine the unique functionality associated with different ceramides within the endothelium^40^. For this study, we utilized C2 ceramide, a cell permeable form of the lipid. However, short chain ceramides do not exist physiologically, and they can be hydrolyzed and re-acytated into other ceramide isoforms^52^, although the exact isoforms formed within the endothelium during health versus disease remain unknown. As such, the dual dependency on both NO and H_2_O_2_ following ceramide administration during NSmase inhibition may not be reflective of the ceramide isoforms naturally formed by NSmase. This may explain why administration of exogenous S1P fully restored NO-dependent dilation during NSmase inhibition, unlike C2-ceramide.

Alternatively, C2-ceramide may have direct effects on pro-oxidative stress pathways^53, 54^. As such, the ceramides that get converted to S1P may increase NO while any remaining C2-ceramides may increase H_2_O_2_. In arterioles from patients with CAD, exogenous C2-ceramide did not fully restore FID during NSmase inhibition. Again, this may reflect differences between endogenously produced ceramide versus C2-cermaide conversion/signaling in diseased states.

Alternatively, it may reflect an important role of sphingomyelin degradation in diseased arterioles during flow. In an ideal scenario, we would like to administer various chain lengths of ceramide during NSmase inhibition, however there are technical challenges in dissolving and administering long-chain fatty acids to isolated arterioles. We would also like to be able to measure sphingolipid content in the microvascular endothelium following various treatments, however nearly 20-30 microvessels per patient would have to be pooled to obtain enough starting material for tandem-mass spectrometry. Even if this approach could be implemented, such measurement would be reflective of all cell types within the vessel and may not be sensitive enough to identify endothelial-specific changes given that smooth muscle cells make up the vast majority of an arteriole.

There are important limitations to this study. While studying human microvascular function is critical for informing translational mechanisms, this approach limits our ability to control for lifestyle, non-cardiac related co-morbidities and medication, and other patient characteristics that may be key influencers of vascular function. We instituted a mechanism of consenting patients prior to surgery to receive information about the above-mentioned factors, however, the tissue that we receive through the discarded tissue bank still has limited information. Nonetheless, we receive information on sex, age, race, CAD risk factors and related to medications on all tissue regardless of mode of collection. This allows us to compare between nonCAD and CAD. To limit the effect of any medications, all vessels are washed for a minimum of 60 min prior to initiation of studies. Additionally, the use of surgical samples necessitate that we group patients with 0 or 1 risk factor of CAD as “healthy.” There are also sex and age differences between nonCAD and CAD patients, which are reflective of the surgical population. However, data suggests that age^24, 55^ or sex^24^ alone do not significantly influence ex-vivo microvascular function. Despite these limitations, the use of human arterioles is crucial for evaluating mechanistic differences during health and disease and informing translational targets for promoting microvascular health.

### Perspectives and Conclusions

While growing evidence highlights microvascular function as a key predictor of and contributor to cardiovascular disease outcomes, there are currently no targeted strategies available for treating or preventing microvascular dysfunction. The well-established detrimental effects of ceramide on cardiovascular function have sparked ample research that aims to decrease ceramides. However, our study highlights that despite key differences in downstream signaling between health and disease, acute NSmase-mediated ceramide formation and its subsequent conversion to S1P is necessary for proper functioning of the human microvascular endothelium. As such, therapeutics that aim to significantly lower ceramide levels, especially through neutral sphingomyelinase inhibition, may actually prove detrimental to the vasculature.

## Acknowledgements

The authors would like to thank Sergey Tarima, PhD from the Division of Biostatistics at the Medical College of Wisconsin for his assistance with statistical analysis.

## SUPPLEMENTAL FIGURES AND TABLES

**Supplemental Figure 1.**
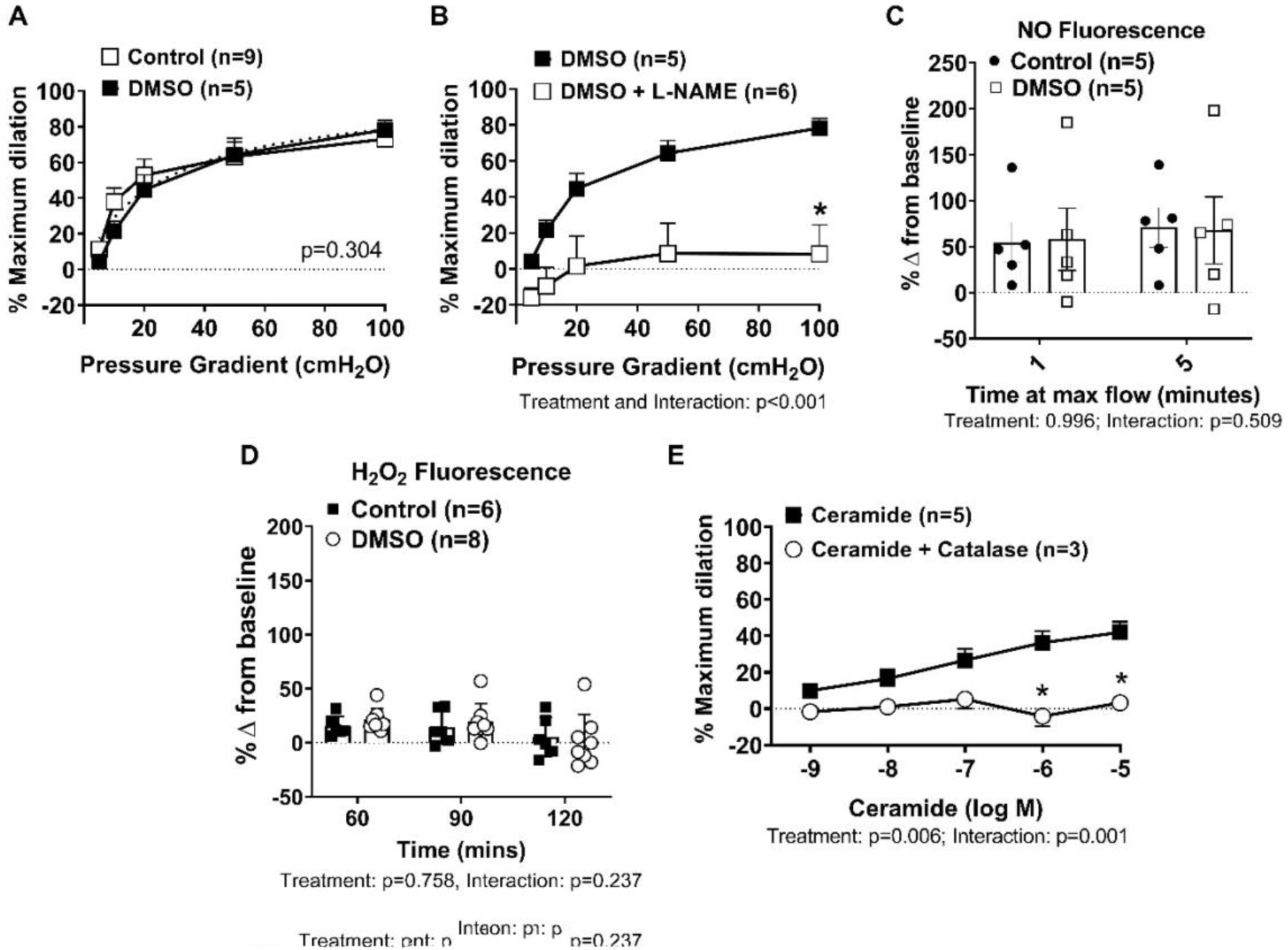
(A) Flow response following vehicle control treatment (DMSO, 30mins, vol/vol; n=5) compared to untreated control (n=9). (B) Flow response with vehicle control (DMSO) treatment in the presence of L-NAME (100μM; n=6) compared to vehicle control alone (n=5; 30mins). (C) NO fluorescence intensity following 1 and 5 minutes of maximal flow (100cmH_2_O) in control (n=5) and vehicle control (n=5) treated human arterioles. (D) H_2_O_2_ fluorescence intensity over time for up to 120 mins in control (n=6) and vehicle control (n=8) treated HUVECs. (E) C2-Ceramide-induced dilation (10^-9^ to 10^-5^ M, 1min; n=5) in the presence of catalase (500U/ml; 30mins; n=3) compared to control (n=5). (A) – non-linear regression with a 4-parameter fit (min constrained to 0, max constrained to 100, and EC50 constrained to >0); dotted line represents best-fit. (B, C, D, E) – two-way repeated measured ANOVA with Holm-Sidaak multiple comparisons test. p<0.05

**Supplemental Figure 2.**
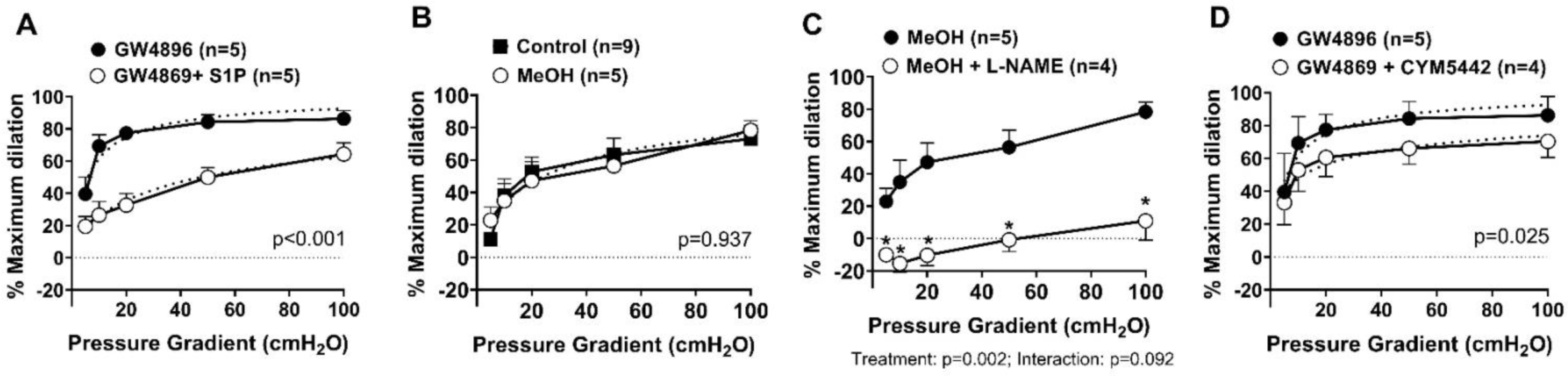
(A) Flow response in vessels treated with S1P (1μM, 30mins) + GW4869 (4μM, 30mins; n=5) compared to GW4869 alone (n=5). Response to flow with vehicle control (MeOH, vol/vol; n=5) (B) compared to control (n=9) and (C) with L-NAME (100μM; n=4). (D) Flow response in vessels treated with an agonist of S1PR1 (CYM5442, 1μM, 30mins) + GW4869 (4μM, 30mins; n=4) compared to GW4869 alone (n=5). (A, B, D) – non-linear regression with a 4-parameter fit (min constrained to 0, max constrained to 100, and EC50 constrained to >0); dotted line represents best-fit. (C) – two-way repeated measured ANOVA with Holm-Sidaak multiple comparisons test. *p<0.05

**Supplemental Figure 3.**
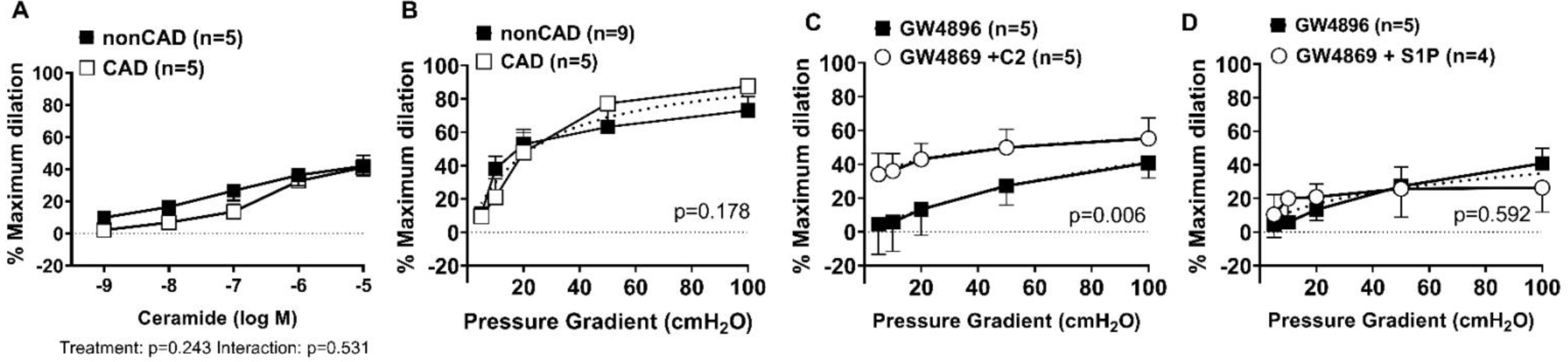
(A) Exogenous C2-ceramide induced dilation (10^-9^ to 10^-5^ M; 1min) of arterioles from patients with CAD (n=5) compared to arterioles from healthy, nonCAD patients (n=5). (B) Flow response of arterioles from patients with CAD (n=5) compared to arterioles from healthy, nonCAD patients (n=9). Flow response of CAD arterioles with (C) exogenous ceramide (10μM) + GW4869 (4μM, 30mins, n=5) and (D) exogenous S1P (1μM) + GW4869 (n=4) compared to GW4869 alone (n=5). (A) – two-way repeated measured ANOVA with Holm-Sidaak multiple comparisons test. (B-D) – non-linear regression with a 4-parameter fit (min constrained to 0, max constrained to 100, and EC50 constrained to >0); dotted line represents best-fit.

**Supplemental Table 1.** Patient Demographic Information. *Two-tailed t-test ^#^Chi-square analysis

|  | Healthy (nonCAD;<br>n=91) | CAD (n=32) | p-value |
| --- | --- | --- | --- |
| Age (mean±SD) | 44.2±12.3 | 65.3±9.7 | <0.001* |
| Sex | 83 female, 8 male | 12 female, 20 male | <0.001# |
| Race/ethnicity | 67 Caucasian, 13 African American, 4 Hispanic, 1 Asian, 4 unknown/Other | 27 Caucasian, 4 African American, 1 unknown/Other | 0.867# |
| BMI (mean±SD) | 32.2±8.3 | 30.1±5.5 | 0.186* |
| Hypertension | 6 | 25 | <0.001# |
| Congestive heart failure | 0 | 8 | <0.001# |
| Myocardial infarction | 0 | 10 | <0.001# |
| Stroke | 0 | 1 | 0.09# |
| Peripheral vascular disease | 0 | 5 | <0.001# |
| Diabetes | 0 | 11 | <0.001# |
| Hyperlipidemia | 5 | 21 | <0.001# |
| Smoker | 1 | 5 | 0.001# |
| Cancer | 4 | 5 | 0.036# |

**Supplemental Table 2.**
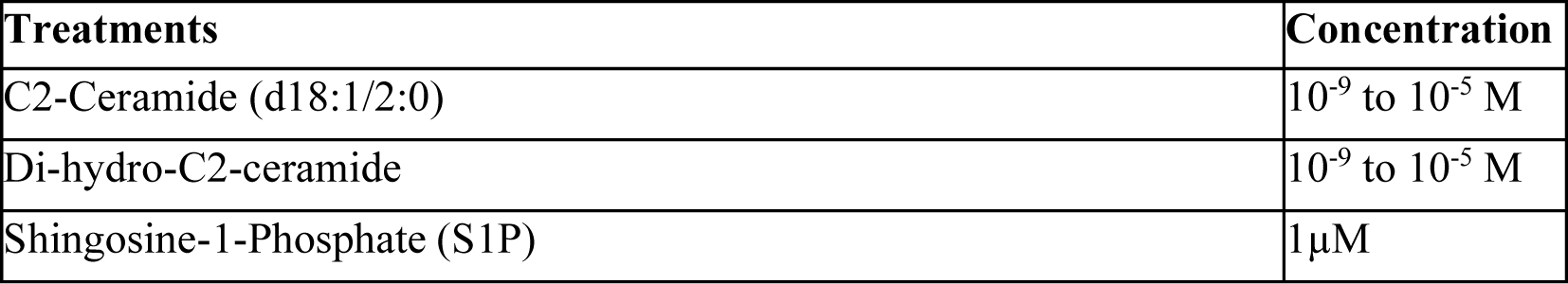
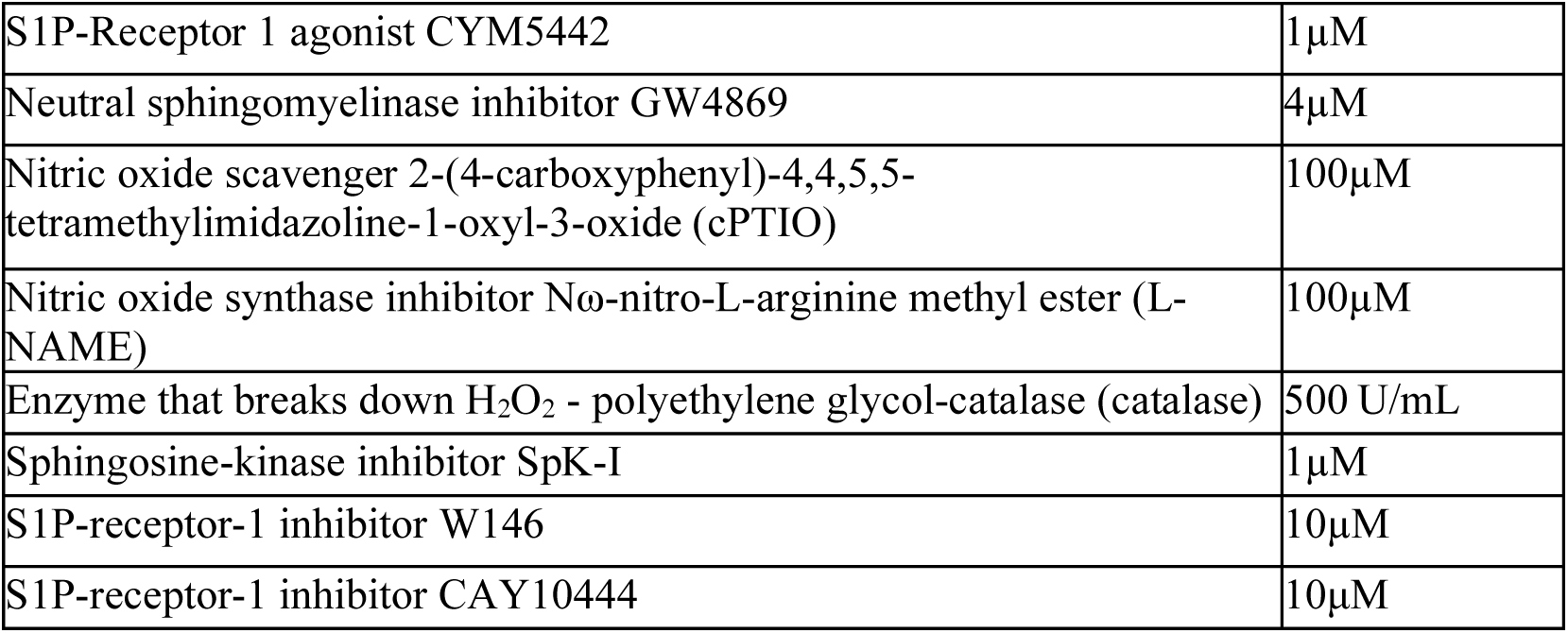
Treatment Details

| Treatments | Concentration |
| --- | --- |
| C2-Ceramide (d18:1/2:0) | $10^{-9}$ to $10^{-5}$ M |
| Di-hydro-C2-ceramide | $10^{-9}$ to $10^{-5}$ M |
| Shingosine-1-Phosphate (S1P) | $1\mu\text{M}$ |
| S1P-Receptor 1 agonist CYM5442 | 1μM |
| Neutral sphingomyelinase inhibitor GW4869 | 4μM |
| Nitric oxide scavenger 2-(4-carboxyphenyl)-4,4,5,5-tetramethylimidazoline-1-oxyl-3-oxide (cPTIO) | 100μM |
| Nitric oxide synthase inhibitor Nω-nitro-L-arginine methyl ester (L-NAME) | 100μM |
| Enzyme that breaks down H<sub>2</sub>O<sub>2</sub> - polyethylene glycol-catalase (catalase) | 500 U/mL |
| Sphingosine-kinase inhibitor SpK-I | 1μM |
| S1P-receptor-1 inhibitor W146 | 10μM |
| S1P-receptor-1 inhibitor CAY10444 | 10μM |

